# Accelerated maturation of branched organoids confined in collagen droplets

**DOI:** 10.1101/2025.03.24.644794

**Authors:** Iris Ruider, Anna Pastucha, Marion K. Raich, Wentao Xu, Yan Liu, Maximilian Reichert, David Weitz, Andreas R. Bausch

## Abstract

Droplet-based organoid culture offers several advantages over conventional bulk organoid culture, such as improved yield, reproducibility, and throughput. However, organoids grown in droplets typically display only a spherical geometry and lack the intricate structural complexity found in native tissue. By incorporating singularized pancreatic ductal adenocarcinoma cells into collagen droplets, we achieve the growth of branched structures, indicating a more complex interaction with the surrounding hydrogel. A comparison of organoid growth in droplets of different diameters showed that while geometrical confinement improves organoid homogeneity, it also impairs the formation of more complex organoid morphologies. Thus, only in 750 *µ*m diameter collagen droplets did we achieve the consistent growth of highly branched structures with a morphology closely resembling the structural complexity achieved in traditional bulk organoid culture. Moreover, our analysis of organoid morphology and transcriptomic data suggests an accelerated maturation of organoids cultured in collagen droplets, highlighting a shift in developmental timing compared to traditional systems.

## Introduction

Progenitor cells isolated from human adult tissue or stem cells are able to navigate a complex spatiotemporal differentiation pattern that gives rise to organoid structures that capture important properties of native tissue [1–4]. Many developmental processes result in branched structures as found in several distinct tissues, including the lungs [5, 6], kidneys [7, 8], mammary glands [9, 10] and pancreas [11, 12]. This process of branching morphogenesis also plays a relevant role in pathological conditions such as tumor progression in pancreatic ductal adenocarcinoma (PDAC). Indeed, PDAC organoid growth successfully replicates the various developmental processes and maturation steps characteristic of PDAC tumors in vivo [13]. Beyond scalability and parallelization, drug-screening applications rely heavily on the ability to steer and control such maturation steps [14]. The use of droplet microfluidics [15, 16] to encapsulate cells into small hydrogel droplets or core-shell structures has enhanced the throughput [17, 18] and scalability [19] of organoid culture systems. Most of those recent applications focused on establishing spheroid culture systems from cell culture aggregates in matrigel without higher-order structure formation processes [20–26]. Single-cell-derived organoids from progenitor cells cultured in hydrogel droplets displaying distinct developmental phases remain to be implemented. Confining organoid culture to small droplets alters the mechanical properties of the surrounding hydrogel by the modified boundary condition. Its impact on the developmental processes during organoid growth remains unexplored. Here, we demonstrate that branching organoid structures can be grown in collagen droplets, recapturing faithfully the structures found in bulk assays and even in vivo. To this end, we incorporated singularized murine pancreatic ductal adenocarcinoma cells into the collagen droplets and tightly controlled the temperature gradient in the microfluidic device, ensuring uniform collagen polymerization within the hydrogel droplets. We found that 750 *µ*m diameter droplets were ideally suited to enable branching morphogenesis and lumen formation in single-cell derived PDAC organoids. We showed that geometrical confinement accelerates organoid maturation, further highlighting the potential of this approach for high throughput and time-sensitive experiments.

### Generation of collagen droplets using droplet-based microfluidic chips

To generate collagen droplets, we used PDMS microfluidic chips with two inlets: one for the collagen solution mixed with singularized PDAC cells and another one for the fluorinated oil with surfactant (Fig. 1 a and Extended Data Fig. 1 a). Collagen droplets were generated at the microfluidic chip’s cross-section and collected in a heated reservoir at 37 °C. To achieve full collagen polymerization, the droplets were maintained in emulsion at 37 °C for 1 hour (Fig. 1 a). Subsequently, the droplets were recovered from the emulsion and transferred to cell culture medium. The cells were cultured in the collagen droplets for up to a week and developed into branched organoids (Fig. 1 a). The microfluidic chip was operated in a cold room to prevent premature collagen polymerization before droplet formation. We performed experiments to assess the impact of the microfluidic setup and the collecting reservoir temperature on collagen polymerization (Fig. 1 b). Our findings indicate that polymerization at 37 °C ensures evenly polymerized hydrogel (Fig. 1 c, d and e). Moreover, it is crucial to maintain the temperature of the collagen at 4 °C throughout the seeding process to prevent premature collagen polymerization (Fig. 1 c and d). The collagen polymerizes homogeneously only under such tightly controlled temperature gradients (Fig. 1 e). To investigate the impact of droplet size on organoid growth behavior, we fabricated microfluidic devices of two different heights, producing collagen droplets of 373± 21 *µm* (small droplets, mean± s.d.) and 756± 30 *µm* (large droplets, mean ± s.d.) in diameter (Extended Data Fig. 1 b). The production rates were 10 Hz for the small droplets and 1 Hz for large droplets. A cell in a large droplet experienced a microenvironment with 8 times the collagen volume compared to a cell in a small droplet. The collagen solution containing the singularized PDAC cells was well-mixed to ensure even cell distribution in the seeding volume. H2B-Teal expressing PDAC cells were encapsulated in hydrogel spheres following a Poisson distribution during droplet formation. Consequently, 37% of large droplets were empty, 36% contained a single cell, and 27% contained two or more cells. For small droplets, 33% were empty, 33% contained one cell, and 34% contained two or more cells (Extended Data Fig. 1 c). We determined the proportion of droplets containing at least one developed three-dimensional structure on day 4 to quantify organoid growth efficiency in the hydrogel spheres. For large droplets, containing on average 1.1 cells per droplet, 47 ± 5% (mean± s.d.) of droplets contained at least one organoid on day 4. small droplets, containing on average 0.8 cells per droplet, showed a slightly lower efficiency, with 40± 8% (mean ± s.d.) of droplets containing at least one organoid on day 4 (Extended Data Fig. 2). Thus, droplet-based organoid culture yielded approximately one organoid for every second seeded cell, which represents a significantly higher efficiency than compared to bulk, where extreme limiting dilution analysis indicates that only every fifth cell forms an organoid [13].

**FIG. 1.**
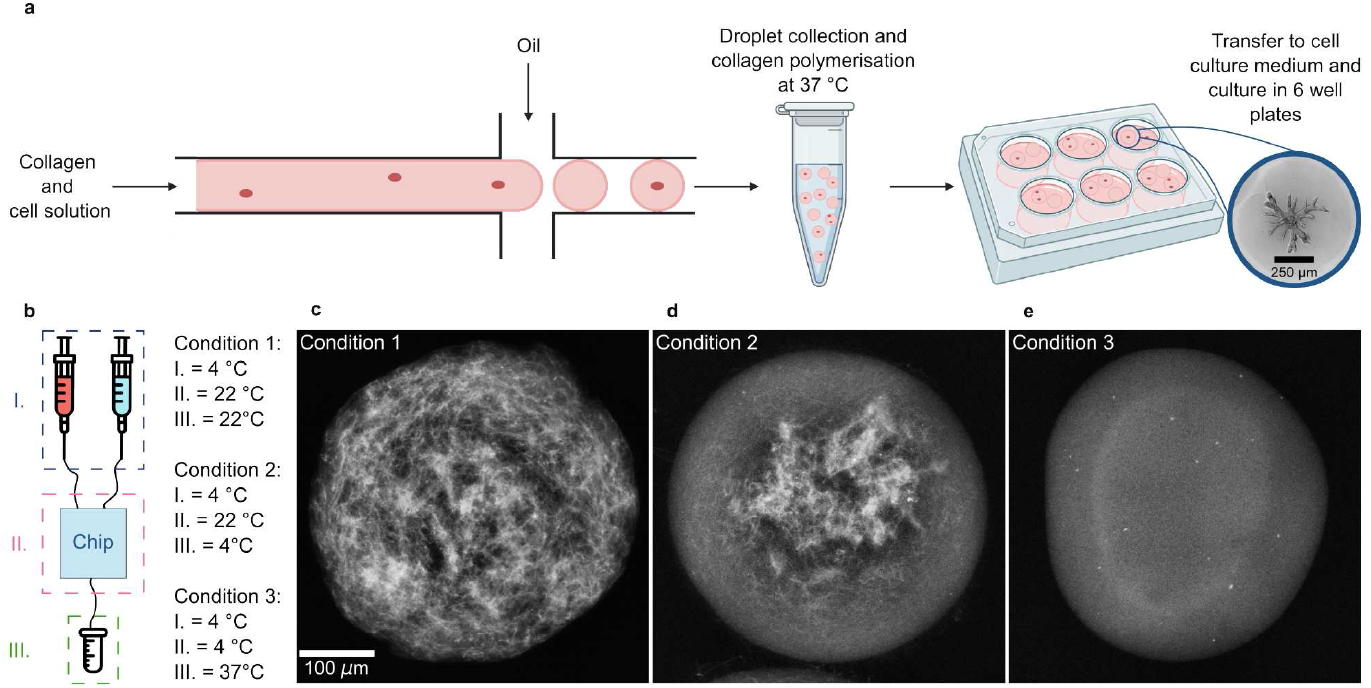
Microfluidic setup for organoid production. (a) Singularized cells are encapsulated in collagen droplets using droplet based microfluidics. After breaking the emulsion, the droplets are transferred to cell culture medium, and the organoids are cultured for 8 days. In large droplets, the seeded cells develop into branched organoids. (b) The microfluidic chip was operated with three different temperature gradients throughout the pipeline to establish optimal collagen polymerization conditions. (c) At room temperature, the collagen droplets polymerize heterogeneously because of collagen fiber aggregation (n = 2 independent experiments). (d) The microfluidic chip is operated at room temperature, and the reservoir is cooled during droplet collection. Afterward, polymerization is induced at 37 °C. The collagen polymerized prematurely as the temperature control was disrupted during droplet production (n = 2 independent experiments). (e) Maintaining the whole setup at 4 °C and the collecting reservoir at 37 °C allowed for homogenous collagen polymerization throughout the droplet (n = 2 independent experiments).

**FIG. 2.**
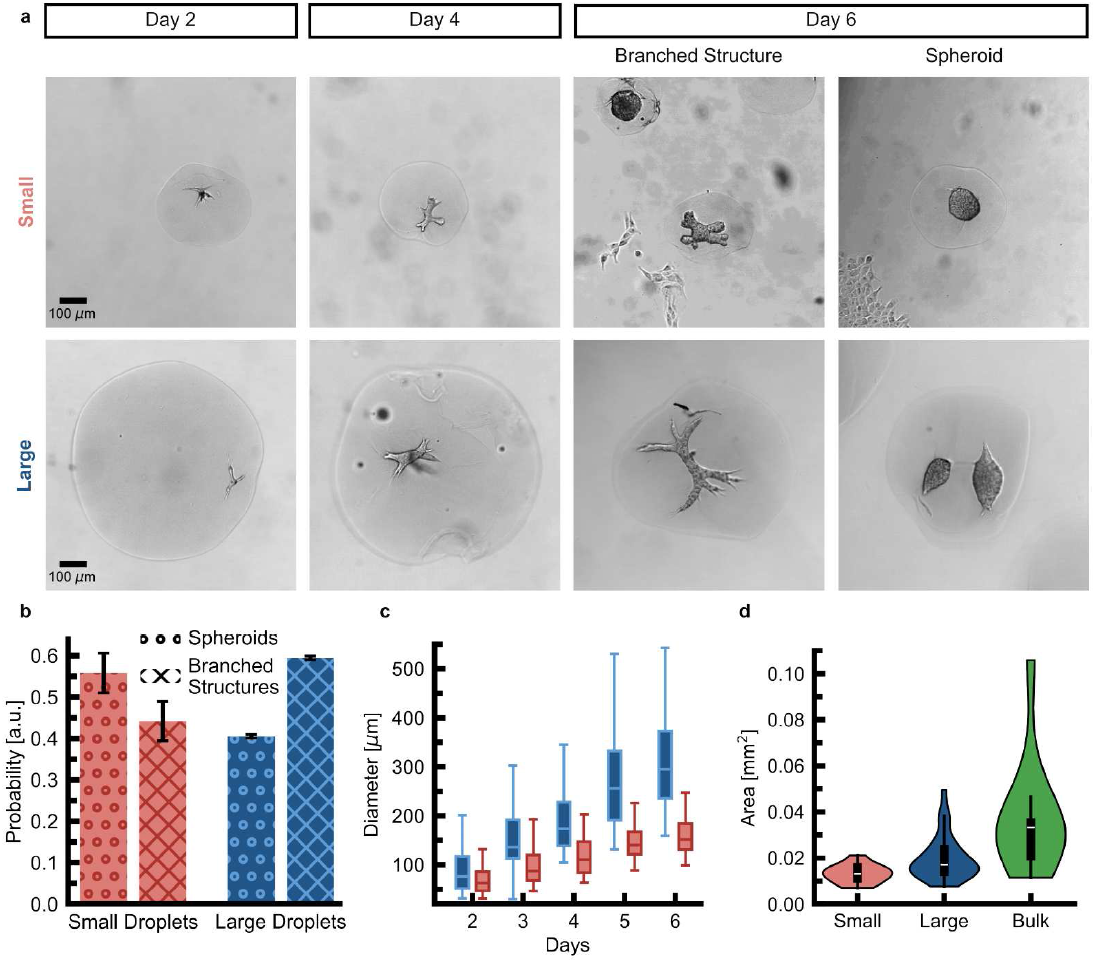
Characterization of organoid morphology in _Small_ versus large droplets. (a) Morphology of organoids grown in large and _Small_ droplets over time (n_Large_ ≥ 34 and n_Small_ ≥ 24 for each day). (b) Probability of obtaining spheroids versus branched structures in _Small_ (red) versus large (blue) droplets (N = 3 independent experiments with n_Small_ ≥ 116 and n_Large_ ≥ 153 structures evaluated for each experiment). Error bars indicate the standard deviation. (c) The major axis measured for organoids grown in _Small_ (red) and large (blue) droplets (n_Large_ ≥ 34 and n_Small_ ≥ 24 for each day). (d) The distribution of the area for 2D projected organoids on day 6 for organoid culture in _Small_ (red) and large droplets (blue) and in bulk (green) (n_Small_ = 24, n_Large_ = 60, n_Bluk_ = 13).

### Characterization of organoid growth in small and large collagen droplets

Single PDAC cells, seeded in collagen gels with a volume of approximately 1 ml, develop into mature structures with dimensions reaching several millimeters. By day 5, PDAC organoids typically measure 542± 155 *µm*(mean± s.d.) along their major axis [9, 13]. To investigate the impact of volume confinement, we analyzed organoid development in small(≈ 370 *µm* diameter, 27 nl) and large(≈ 750 *µm* diameter, 221 nl) collagen droplets. Organoids in small droplets had an increased tendency to grow in a spherical shape. In contrast, organoids in large droplets tended to evolve into elongated structures with branched morphologies (Fig. 2 a). We analyzed segmented 2D projections of the organoids and computed their compactness over time by comparing the parameter of the organoid’s shape to that of a circle with the same area. We found that organoids grown in small droplets displayed a lower compactness after 6 days in culture compared to organoids grown in large droplets, supporting the hypothesis that organoids in small droplets rather grew as spheroids (Extended Data Fig. 3). To evaluate the efficiency of branching morphogenesis, we screened large and small droplets four days after seeding and quantified the proportion of spheroids versus branched structures. Given that the first 5 days of bulk organoid culture are characterized as the onset phase, we distinguished between spheroid structures and invasive phenotypes [13]. Spheroids were defined as structures with a major axis *a* inferior to 60 *µm*, a minor axis b greater than a/2, or fewer than 2 branches of at least 30 *µm* in length. Structures that did not meet these criteria were classified as branched, displaying at least 2 branches or an invasive phenotype with the potential for further branching in later developmental stages. Our analysis revealed that in small droplets, only 44± 5% (mean ± s.d.) of the structures were classified as branched structures, whereas in large droplets, 59 ± 5% (mean ± s.d.) did fall under this classification (Fig. 2 b). The major axis of organoids in large droplets increased more rapidly than in small droplets (Fig. 2 c). By day 5, organoids in small droplets measured 150± 42 *µm* (mean± s.d.) along their major axis, while those in large droplets reached a mean major axis of 266 ± 89 *µm* (mean± s.d.). Despite this, even organoids in large droplets only attained half the size of those grown in bulk organoid gels. To assess if droplet-grown organoids displayed a more homogeneous size distribution, we measured the area of the 2D projected organoids, A_pro_j, and computed the polydispersity index 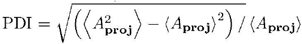 as an estimate of the homogeneity of the organoid culture [18]. The polydispersity index for bulk-grown organoids is *PDI=* 63%. The one for organoids grown in large droplets decreases to *PDI* = 47%. The polydispersity index for organoids grown in small droplets is reduced to *PDI* = 28%. Therefore, a restriction in the available volume of the hydrogel improves the homogeneity of the organoid culture (Fig. 2 d), while at the same time reducing morphological complexity.

**FIG. 3.**
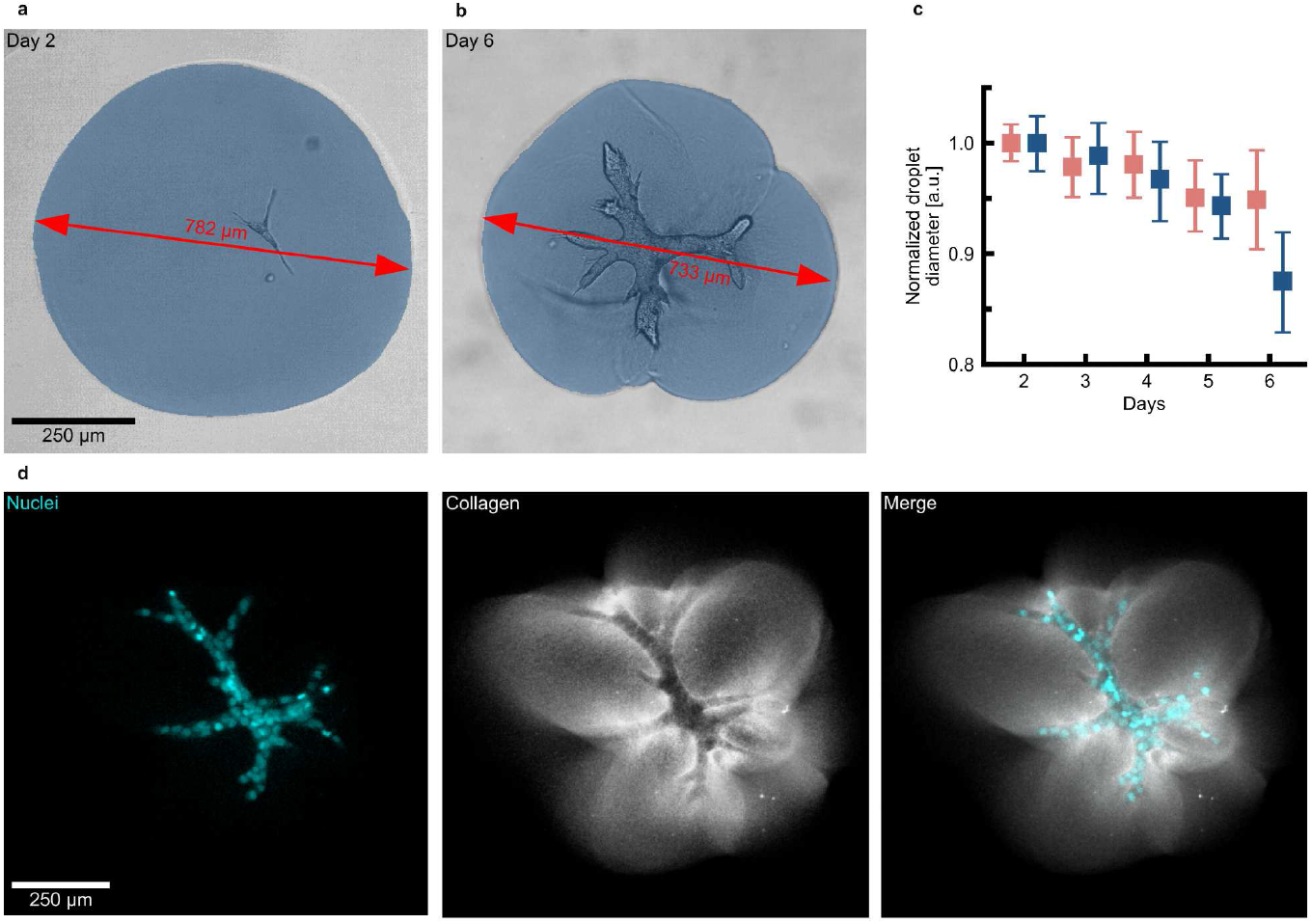
Organoids compress the collagen droplets. (a) The droplets’ surface is homogenous 2 days after organoid seeding as the growing organoid does not yet exert sufficient mechanical force to deform the collagen matrix (n = 32 droplets). The segmented collagen droplet is visualized in blue. (b) The droplet’s surface is increasingly irregular 6 days after organoid seeding, as the branched organoid pulls on the surrounding matrix and causes increasing droplet deformation, resulting in a reduced diameter (n = 29 droplets). (c) For large droplets, branched structures cause a 12% reduction in droplet diameter because of collagen deformation. In contrast, small droplets only undergo 5% of diameter reduction on day 6 (n_Large_ ≥ 23 and n_Small_ ≥ 23 for each time point). Error bars indicate the 95% confidence interval. (e) The deformation of a collagen droplet by an organoid cultivated for 5 days leads to the formation of a collagen cage. The collagen deformation is the highest at the tips of the organoid’s branches (n = 2 independent experiments).

### Characterization of collagen interaction

The formation of branched structures depends on the cells’ ability to interact and plastically deform their surrounding matrix [9, 13]. By pulling on the collagen fibers, the organoid forms a collagen cage that facilitates the development and growth of branched structures. Given the increased branching ten dency of organoids in large droplets, we investigated if these organoids also displayed increased interaction with their surrounding matrix. To assess this, we monitored changes in droplet diameter over time. _Large_ droplets underwent visible deformation after one week in culture (Fig. 3 a and b). We measured the 2D-projected shape of the droplets to quantify these deformations and examined whether the continuous pulling exerted by the organoids led to a decrease in droplet size. Our results showed that small and large droplets experienced similar amounts of compression up to day 5. However, by day 6, large droplets were compressed by 12%, compared to only 5% for small droplets (Fig. 3 c). Next, we compared organoid morphology and droplet deformation. Assuming the organoids grew in the center of the droplets, we calculated the relative distance between the organoid and the microsphere boundary, defined as (r-a)/r. Up to day 5, the organoids in small droplets were closer to the boundary than those in large droplets (Extended Data Fig. 4). However, by day 6, the organoids in large droplets were closer to the outer boundary than those in small droplets, suggesting more extensive growth and mechanical deformation in larger droplets at day 6. To examine how complex morphologies, such as branched structures, influence mechanical deformation, we used fluorescently labeled collagen and stained PDAC cell nuclei with SiRDNA to visualize the interaction between the branched structures and the collagen matrix. By day 5, branched structures were observed to deform the surrounding collagen. Indeed, the areas with the most deformation coincided with the locations of organoid branches. Additionally, the increased collagen intensity at the organoid boundary suggested the formation of a collagen cage, which is crucial for the cells to expand into a branched structure (Fig. 3 e).

**FIG. 4.**
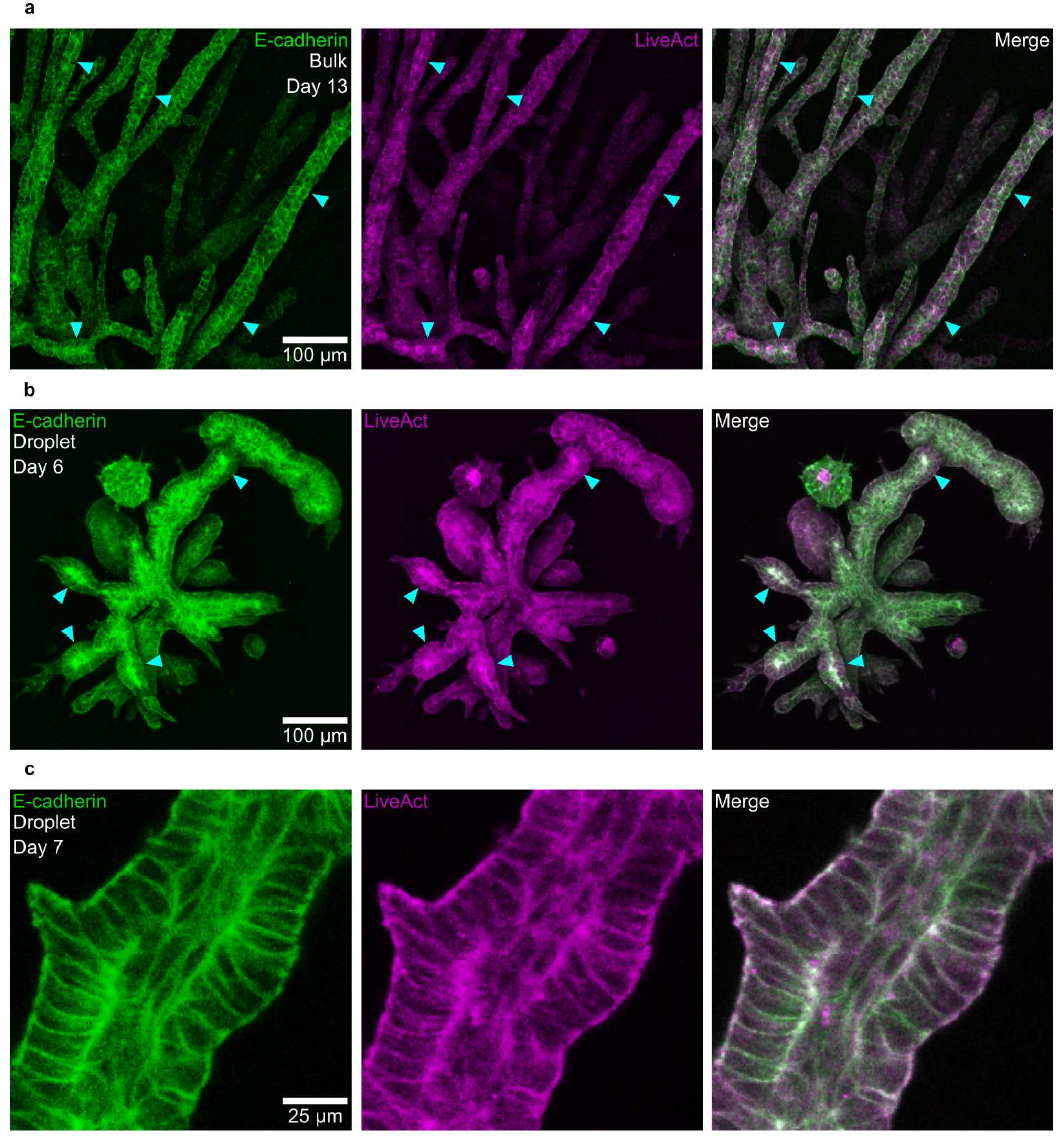
Branched Organoids in large droplets display similar developmental stages to those in bulk. (a) PDAC organoids cultured in bulk display local heterogeneous expression levels of E-Cadherin and F-actin that indicate lumen formation sites on day 13 (n = 3 independent experiments). (b) PDAC organoids grown in large droplets display local heterogeneous expression levels of E-Cadherin and F-actin, indicating lumen formation sites already on day 6 (n = 3 independent experiments). (c) A fully formed lumen in a branched organoid grown in a large droplet on day 7 (n = 3 independent experiments).

### Characterization of organoid morphology

In bulk, branched PDAC organoids exhibit a characteristic lumen formation phase in their final development stage [13]. Leading up to lumen formation, the organoids undergo increasing epithelialization, which is marked by a buildup of E-Cadherin and F-actin at the future apical sites where the lumen will form (Fig. 4 a). Lumen formation sites can be observed in bulk-grown organoids as early as day 11 [27]. In contrast, droplet-grown organoids display a notably faster progression. On day 4, the droplet-grown organoids still display a mesenchymal phenotype characteristic for bulk-grown organoids on day 7 (Extended Data Fig. 5 a and b). However, by day 6, clear signs of epithelialization are already evident, with E-Cadherin and F-actin accumulation at lumen nucleation sites within the branches, along with some fully opened lumens on day 7 (Fig. 4 b and c). This accelerated development in droplet-grown organoids results in lumen formation and maturation occurring six days earlier than in bulk-grown organoids, indicating a more rapid maturation process.

**FIG. 5.**
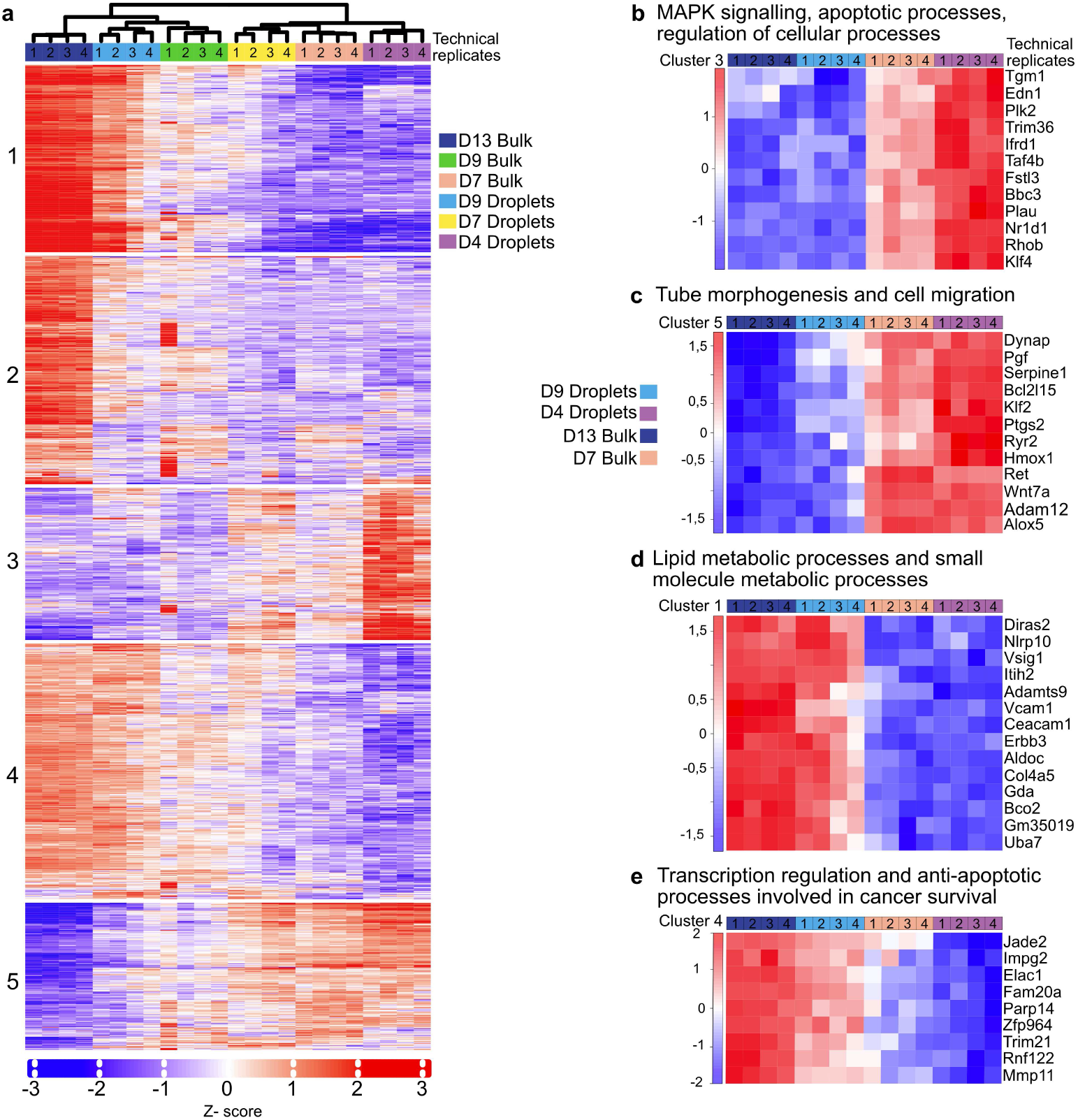
PDAC organoids expression pattern in different growth conditions. (a) K-Means clustering with five clusters (k = 5) using 2000 genes with the most variable expression level between different growth conditions: droplets day 4, 7, 9 and bulk day 7, 9, 13 (n = 4 individual experiments), FDR = 0.05. (b) GO Biological Processes enrichment analysis from cluster 3 for selected conditions (Droplets day 4, 9 and _Bluk_ day 7, 13) highlights genes involved in MAPK signaling and apoptosis-related processes. (c) Gene expression heatmap (cluster 2) for selected genes regulating tube morphogenesis and cell migration. (d) Gene expression heatmap (cluster 1) of a subset of genes involved in lipid metabolism and small molecule metabolism. (e) Heat map for selected genes (cluster 4) involved in transcription regulation and anti-apoptotic pathways facilitating cancer survival.

### Characterization of the transcriptome for organoids grown in large droplets versus bulk

Our imaging data showed that organoids grown in large droplets developed key morphological features and reached developmental hallmarks five days earlier than those in bulk organoid culture. To further explore this, we compared the transcriptomes of organoids grown in bulk culture to those grown in large droplets. As previously reported, bulk organoid culture shows a stage-specific gene expression pattern, with genes being up- or downregulated according to the developmental phase. Early-stage organoids exhibit a mesenchymal-like phenotype, transitioning to a more epithelial-like phenotype at later stages [13]. To evaluate how the geometrical confinement effects influence these gene expression dynamics, we analyzed the top 200 most variably expressed genes across all samples, using Euclidean clustering to visualize a gene expression heat map. In bulk culture, we observed the expected oscillatory pattern, where certain genes were downregulated at day 7 and upregulated at day 13 as the organoids matured (Fig. 5 a). Day 9 represented a transition point, with partially overlapping gene expression profiles between the two states. In droplet-grown organoids, the gene expression profile on day 9 closely resembled the profile on day 13 in bulk culture, consistent with the accelerated maturation timeline observed in their morphology. Given this shift in the timing of gene regulation, we conducted a deeper analysis of the expression patterns across different gene clusters. We identified a switch in the gene expression pattern involved in critical metabolic processes, such as lumen formation and branching morphogenesis, relevant during PDAC organoid maturation (Fig. 5 b and c) [13, 27]. For later time points (D13 bulk and D9 droplets), we detected a population of overexpressed genes involved in cancer survival and lipid metabolic processes characteristic for enhanced metabolic activity during tumor progression (Fig. 5 d) [28]. In droplet-based organoid culture at day 9, genes responsible for cancer survival and anti-apoptotic processes follow the characteristic upregulation pattern observed in bulk organoid culture at day 13, indicating faster maturation (Fig. 5 e).

## Discussion

Encapsulating singularized cells into collagen droplets resulted in the formation of individual highly branched organoid structures. Thus, droplet-based organoid culture improved yield and sample homogeneity and achieved a faster maturation of the organoids. A reduction in droplet size, while improving organoid homogeneity, entailed a loss in morphological complexity. Thus, only in large droplets did we obtain organoids with similar structural complexity to organoids grown in bulk culture. Moreover, it demonstrates the feasibility of cultivating droplet-grown organoids with complex morphologies rather than spherical shapes. Using microscopic and transcriptomic analysis, we confirmed the accelerated maturation of organoids grown in geometrically confined microenvironments, which aligns with previous studies on stem cell aggregates cultured in microwells [29]. Our findings demonstrate that mechanical boundary conditions are fundamental for steering organoid structure formation processes. This highlights the importance of understanding how altered geometrical and mechanical microenvironments affect maturation timelines. Although the mechanisms behind the observed accelerated maturation remain unclear, our findings underscore the need to account for potential shifts in timescales when using droplet-based organoid cultures. Such shifts may impact critical aspects of dose-response studies, including the timing of drug exposure, sensitivity to treatments, and the reproducibility of results.

## ACKNOWLEDGEMENTS

The primary tumor cells (PDAC) Cdh1-mNeonGreen cell line is a kind gift from Prof. Dieter Saur’s laboratory. The Phoenix ECO helper-free retrovirus producer cell line and the Teal-H2B plasmid (#24_pLNCX2_H2B-Te) are a kind gift from the laboratory of Prof. Carsten Grashoff (Universität Münster). pTK93_Lifeact-mCherry was a gift from Iain Cheeseman (Addgene plasmid # 46357 ; http://n2t.net/addgene:46357; RRID:Addgene_46357). We gratefully acknowledge financial support from the European Research Council (ERC) under the European Union’s Horizon 2020 research and innovation program (grant agreement no. 810104-PoInt). This research was conducted within the Max Planck School Matter to Life, supported by the German Federal Ministry of Education and Research in collaboration with the Max Planck Society. We acknowledge partial funding from the Wyss Institute.

## AUTHOR CONTRIBUTIONS

I.R. performed experiments with A.P.’s support under the supervision of A.R.B. M.K.R. generated the cell lines used in this study and performed bulk-organoid experiments. I.R and W.X. and Y.L. conceived the microfluidic chip design and fabricated the masters. A.R.B., and D.W. conceived the project with support of M.R. Data Analysis was performed by I.R. and A.P. All authors participated in the writing of the manuscript.

## COMPETING INTERESTS

The authors declare no competing interests.

## I. MATERIALS AND METHODS

### Provenance of cell lines

Primary tumour cells (PDAC) were gifted by the laboratories of Prof. Maximilian Reichert and Prof. Dieter Saur (Technical University of Munich). Cells were authenticated by genotyping. The Phoenix ECO helper-free retrovirus producer cell line was a gift by the laboratory of Prof. Carsten Grashoff (Universität Münster). All cells were tested and confirmed negative for mycoplasma contamination every 6 months.

### 2D cell culture

Pancreatic ductal adenocarcinoma cells (PDAC) were cultured in high glucose DEMEM medium with 1% of Penicillin/Streptomycin (Thermo Fisher Scientific) and 10% Fetal Bovine Serum (FBS) (Sigma). Cells were grown at 37 °C with 5% of CO_2_ and 80% humidity. Media was exchanged every 48 hours, and cells were passaged by trypsinization upon 80–85% of confluency. For 2D analysis, cell culture with 80-85% confluency was used. Phoenix ECO helper-free retrovirus producer cell line (Phoenix ECO cells) were cultured with high glucose DMEM (Thermo Fisher Scientific) supplemented with 10% FBS and 1% Penicillin-Streptomycin (Thermo Fisher Scientific) in an incubator with an 80% humidity and 5% CO_2_ atmosphere, at 37 °C.

### Generation of fluorescently labelled cell lines

As already described previously [27], for the stable expression of fluorescent markers, retroviral transduction as well as CRISPR/Cas9 transfections were performed with the same primary tumour cells. Retroviral transfection of PDAC cells was implemented during an 8 day protocol using a retroviral plasmid for mCherry-LifeAct (Addgene) [30] or Teal-H2B (gifted by the laboratory of Carsten Grashoff). At day 1, Phoenix ECO cells were seeded in a 175 cm^2^ flasks. Having reached a confluency of 50 to 60% (day 2), Phoenix ECO cells were transfected using Mirus TransIT-X2 Dynamic Delivery System (VWR MIRUMIR-6000) as described in the manufacturer protocol. Media exchange was performed after 24 hours (day 3) of incubation. Simultaneously, PDAC cells were seeded in a 75 cm^2^ flask. Virus was harvested after 48 hours (day 4) from Phoenix ECO cells and sterile filtered (0.45 *µ*m pore size). Supplemented with 7.5 *µ*g/ml polybrene (Sigma-Aldrich) the virus conditioned media was added to the PDAC cells and incubated for 24 hours. This was repeated on day 5. After 48 hours of incubation with virus conditioned media, media was exchanged for PDAC cells with fresh high glucose DMEM (Thermo Fisher Scientific) supplemented with 10% FBS (Sigma F7524) and 1% Penicillin-Streptomyosin (Thermo Fisher Scientific). After 72 hours cells were passaged as described above. Selection of fluorescently labeled cells was implemented by antibiotic resistance, using Puromyocin Dihydrochloride (Gibco) for mCherry-LifeAct, and Geneticin Selective Antibiotic (G418 Sulfate) (Gibco) for Teal-H2B. The FACS sort was carried out using BD Aria Fusion. A CRISPR/Cas9 system was used to endogeneously label E-cadherin in the used primary tumor cells.

### Organoids preparation in bulk collagen

For organoids growing in bulk collagen, we used a previously published protocol [27, 31]. Briefly, PDAC cells (9591) with confluency 80-85% were detached using trypsin and prepared to a final concentration of 500 cells/ml of DEMEM media. Next, the cell suspension was mixed gently with collagen type I solution (rat tail from Corning, NaOH to stabilize pH and neutralizing buffer). The solution was incubated for 1h at 37 °C to polymerize the collagen fibers, and DMEM media was added after detaching the gels from the dish walls. Media was changed first after 72 hours and then every 48 hours. All conditions were performed simultaneously in 4 technical replicates n = 4.

### Chip design

Two different chips were used to achieve different droplet diameters for droplet production. To generate large droplets of 750 *µ*m diameter, a microfluidic chip with a channel height of 600 *µ*m was used, whereas for the production of the smaller sized droplets with a diameter of 370 *µ*m, a chip with 200 *µ*m channel height was used. For both chips, the same underlying CorelCAD design was employed (Extended Data Fig. 1 a). To generate the masters standard soft lithography was performed to fabricate SU8 (Kayaku Advanced Materials, United States) coated silicon wafers (Nano Quarz Wafer, Germany) of 600 *µ*m and 200 *µ*m thickness. The silicon wafers were baked first at 65 °C for 5 minutes and at 95 °C for 40 minutes for 200 *µ*m height and, respectively, for 130 minutes for 600 *µ*m height of SU8. Then, to crosslink the exposed pattern, the SU8-coated wafers were exposed to UV light at an exposure energy of 720 mJ/cm^2^ for 200 *µ*m thick wafers and to 311 mJ/cm^2^ for 600 *µ*m thick SU8 wafers. After UV exposure, the wafers were baked at 65 °C for 5 minutes and at 95 °C for 13 minutes (200 *µ*m thickness) and 33 minutes (600 *µ*m thickness) during the post-exposure bake. The wafers were immersed in SU8 developing solution (PGMEA, Sigma-Aldrich) to develop the structures for 15 minutes (200 *µ*m thickness) and 35 minutes (600 *µ*m thickness) and washed with isopropanol. After developing the structures, the samples were baked at 150 °C for 5 minutes during the post-exposure bake. After master fabrication, PDMS was mixed with curing agent at a ratio of 10:1 (Sylgard 182 Silicone Elastomer, Dow Corning, United States) and poured on the master. To remove air bubbles, the master was then put under vacuum for 15 minutes. Then, the PDMS was cured by overnight incubation in the oven at 70 °C. The next day, the PDMS replica was peeled off the mold. The channel inlets and outlets were generated by using a biopsy puncher of 1 mm diameter. The PDMS was bonded to a glass coverslip after treatment with oxygen plasma. After bonding, the microfluidic chip was incubated at 70 °C for at least 1 hour. Eventually, the microfluidic chips were treated with commercially available Aquapel for 5 minutes at room temperature to obtain a hydrophobic channel surface. The chips were then air dried and incubated at 70 °C for 15 minutes.

### Cell preparation for organoid seeding on the microfluidic chip

The collagen solution for organoid seeding on the microfluidic chip was prepared as previously described for bulk organoid culture [9, 31]. To achieve the incorporation of single cells into collagen droplets the PDAC cells were mechanically separated after Trypsin treatment by aspiration with a syringe needle (28G, Braun). The remaining doublet cells were removed by using a cell strainer (40 *µ*m, Corning). To generate small droplets with single incorporated cells the PDAC cells were diluted to a final concentration of 30 000 cells/ml in collagen, respectively to obtain single incorporated cells in large droplets the cells were diluted to a final concentration of 5 000 cells/ml.

### Droplet production for organoid culture on the microfluidic chip

For droplet production, fluorinated oil Novec 7500 (Iolitec, Germany) with 3% (w/w) surfactant RAN008 (RAN Biotechnologies, United States) was used to generate the small droplets. For the production of large droplets, we used fluorinated oil with 5% surfactant. We loaded the oil with surfactant into 5 ml BD plastic syringes to initiate droplet production. The collagen solution containing the cells was loaded into either a 1 ml BD glass syringe or a 5 ml BD plastic syringe, depending on the required droplet amount. The syringes were connected to the microfluidic chip using fluid dispenser tips (PT-050-27, Drifton, Denmark) and polyethylene micro tubing (PE/2, Scientific Commodities, United States). The syringes were operated using programmable or computer-operated syringe pumps (KDE Technologies, Harvard Pumps or Landgraf HLL). Typical operating speeds for droplet production were 2000 *µ*l/hl and 1000 *µ*l/h for the oil and collagen to generate small droplets or 1000 *µ*l/h and 1000 *µ*l/h for the oil and collagen to generate large droplets. After droplet production, the droplets were collected in a 2 ml Eppendorf tube at 37 °C to initiate collagen polymerization immediately. After droplet production, the Eppendorf tubes were transferred to an incubator and kept at 37 °C and 5% CO_2_ for 1 hour. After collagen polymerization, the fluorinated oil with surfactant was removed, and the droplets were washed three times with only fluorinated oil. After the last washing step, the fluorinated oil was removed, and fluorinated oil with 4% PFO (v/v) was added to the droplets to break the emulsion. Cell culture medium was added on top of the oil phase and the droplets were left for 5 minutes to transition into the aqueous phase. The droplets were centrifuged using a table top centrifuge for 5 minutes to improve droplet recovery. After centrifugation, the pellet formed by the droplets at the interface of the oil and cell culture medium was aspirated with a pipette, and the droplets were transferred into the cell culture medium. The droplets were distributed in 6 well plates. For microscopy imaging, the droplets were transferred to achieve a final concentration of 750 droplets per well, whereas for RNA sample collection, the desired final concentration was 1500 droplets per well.

### Fluorescent collagen preparation

Collagen was fluorescently labeled as previously described [9]. Briefly, collagen was dialyzed at 4 °C to reach pH 7. Subsequently, collagen was conjugated with Atto 488 by overnight incubation at 4 °C. To remove any non-bound dye, dialysis was performed for 8 hours followed by an overnight dialysis step with acid to avoid any premature polymerization. The fluorescently labeled collagen was stored at 4 °C until further usage.

### SiRDNA staining

For nuclei visualization, PDAC organoids were incubated with SiRDNA (Spirochrome SPY650-DNA SC501) for a minimum of 1 h before imaging at 1–2 *µ*g/ml.

### Microscope analysis

Samples were analysed using scanning confocal microscope Leica Sp5 (PL FLUOTAR 10x/0.30 and HC PL APO 20x/0.70), Leica Mica (PL FLUOTAR 10x/0.32) and epifluorescent microscope Leica Thunder (PL FLUOTAR 10x/0.32).

### Characterization of the droplet diameters

Collagen droplets were segmented using Cell Pose 2.0 running on Google Colab [32]. The diameter was extracted from the binary images using the scikit image package.

### Measuring cellular incorporation frequency into droplets

After droplet generation, droplets in 6 well plates were screened for the amount of encapsulated cells.

### Counting the amount of spheroids and branched structures present among organoid phenotypes

Four days after droplet generation, droplets in 6 well plates were screened for the number of structures that developed into spheroids and for the number of structures that displayed an invasive phenotype with the ability to form branches.

### Droplet and organoid segmentation with feature extraction

Binary masks of 2D projected images of collagen droplets containing PDAC organoids and bulk-grown organoids were analyzed using the selection tool in ImageJ (ver. 1.54f) [33]. Python and the scikit-image package were used to analyze morphological features.

### Preparation of RNA samples for mRNA sequencing

2D PDAC cells were cultivated until 80-85% of confluency, detached, and collected by spinning down (300 g) to obtain cell pellets. Organoids on days 4, 7, 9, and 13 were subjected to collagenase treatment (2.6 mg/ml for 1.3 mg/ml collagen and 6 mg/ml for 3 mg/ml collagen) to remove the organoids from the collagen gel. Next, the solution with organoids was spun down (1200 rpm), and the pellet was gently washed with PBS. Total RNA was extracted from the organoids and 2D samples using RNeasy Mini Kit (Qiagen, Cat. No. 74104) according to the manufacturer’s instructions. All steps were performed in RNAse free environment. Next, RNA quality was determined using Agilent RNA 6000 Nano Assay. Library preparation was performed on samples with RIN ≥ 7 and concentration ≥ 30ng/*µ*l. mRNA sequencing was performed by BMKGENE company using Illumina Novaseq X. 6GB of raw data was obtained for each sample. Data was next analyzed using Galaxy Europe server to get normalized read counts [34].

### Differential gene expression (DEG) analysis

We used the DESeq2 R package from iDEP for differential gene expression analysis [35].

### Statistical analysis

Experiments were performed two to three times independently from each other with similar outcomes. For each set of experiments, at least 5 replicates were measured. 95% confidence intervals were computed via bootstrapping with 1000 bootstrap iterations.

## Notes

### Competing Interest Statement

The authors have declared no competing interest.

