## Extended Data for "Accelerated maturation of branched organoids confined in collagen droplets"

### II. EXTENDED DATA

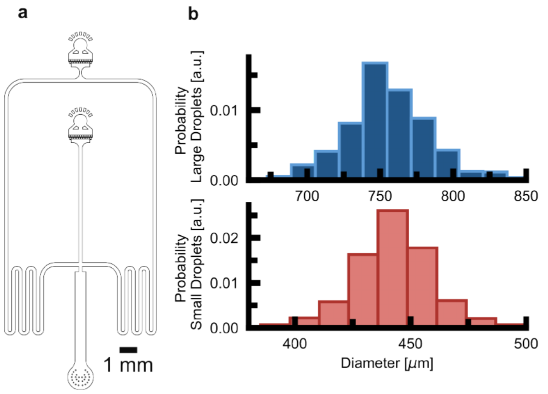


Extended Data Fig. 1. Characterization of the microfluidic setup. (a) The microfluidic chip design. (b) The different heights of the microfluidic devices allow for the generation of large and small droplets, with an average size of 750 $\mu$m and 370 $\mu$m. The histogram for the droplet size distribution is shown in the upper row for large droplets (n = 666 droplets) and in the lower row for small droplets (n = 907 droplets). (c) The incorporation frequency follows Poisson statistics. The cell incorporation frequency for large droplets is displayed in the upper row (n = 951), and the cell incorporation frequency for small droplets is shown in the lower row (n = 906). The line plot represents the theoretical Poisson distribution probabilities, assuming an average of one cell per droplet.

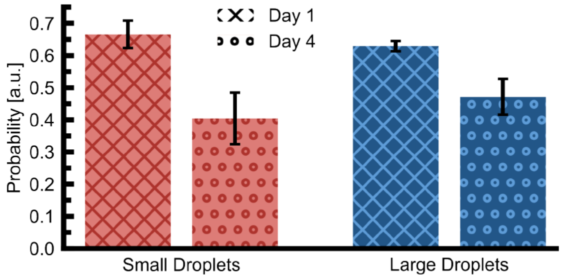


Extended Data Fig. 2. Viability of droplet-grown organoids. Four days after cell seeding, 40 ± 8% of the small droplets (red) contain at least one viable organoid structure compared to 47 ± 5% of the large droplets (blue) (N = 3 independent experiments with $n_{Small} \geq 301$ and $n_{Large} \geq 300$ droplets evaluated for each experiment). The error bars show the standard deviation.

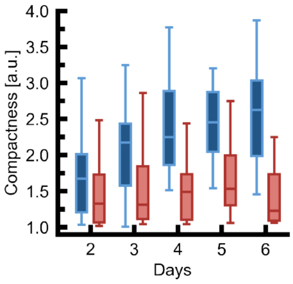


Extended Data Fig. 3. Compactness of 2D-projected organoid shapes. Compactness of the 2D projected shape of the organoids grown in large droplets (blue) and in small droplets (red) over time ($n_{Large} \geq 34$ and $n_{Small} \geq 24$ for each day).

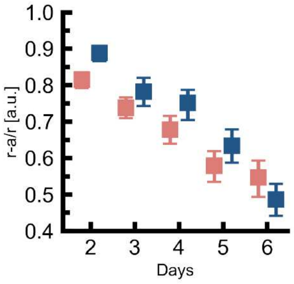


Extended Data Fig. 4. Relative distance between organoid and droplet surface. The mean relative distance between the organoid and the droplets' surface for small droplet (red) and large droplets (blue) over time ($n_{\text{Large}} \geq 23$ and $n_{\text{Small}} \geq 23$ for each time point). The error bars show the 95% confidence interval.

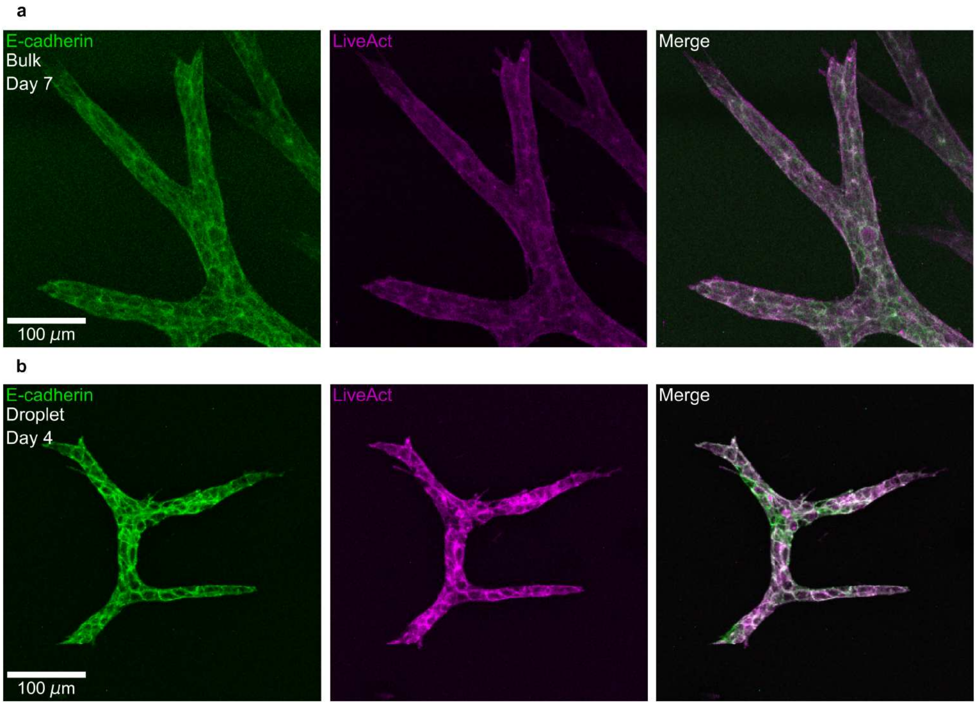


Extended Data Fig. 5. Droplet-grown organoids on day 4 lack epithelialization. (a) The bulk-grown organoid on day 7 displays a homogeneous distribution of E-Cadheerin and F-actin throughout its branches. The cells in the branches display a mesenchymal phenotype in concordance with the organoid's developmental phase (n = 3 independent experiments). (b) The droplet-grown organoid on day 4 shows a homogenous distribution of E-Cadherin and F-actin throughout the branches and displays a mesenchymal phenotype (n = 3 independent experiments).
